# Rubicon modulates neuroimmune responses following traumatic brain injury

**DOI:** 10.64898/2026.03.04.709622

**Authors:** Sagarina Thapa, Amir Mehrabani-Tabari, Olivia Pettyjohn-Robin, Dexter PH Nguyen, Mehari M Weldemariam, Chinmoy Sarkar, Maryam Khan, Maureen A Kane, Marta M Lipinski

## Abstract

Traumatic brain injury (TBI) elicits robust neuroinflammation and oxidative stress, coupled with an acute inhibition of macro-autophagy (autophagy) in neurons and microglia. Rubicon (*Rubcn*), a Beclin1 interacting protein that suppresses autophagy and mediates LC3-associated phagocytosis and endocytosis (LAP/LANDO), influences inflammatory signaling in metabolic, neurodegenerative, and inflammaging diseases; yet its role in acquired brain injury has not been defined. Using a controlled cortical impact model, we investigated the role of Rubicon in acute neuroinflammatory responses following injury by comparing wild-type and *Rubcn*-mutant mice. Bulk-RNA sequencing of injured cortex revealed attenuated induction of inflammatory pathways and reduced activation of pro-inflammatory microglial/macrophage phenotype in injured *Rubcn*-mutant mice. *Rubcn*-mutant mice demonstrated less pronounced inhibition of autophagy during the acute phase of injury. Although the inflammatory differences were transient, Rubicon mutant mice exhibited improved motor coordination and gait stability during recovery. Proteomic analyses revealed the presence of a truncated Rubicon protein in the mutant mice and identified the negative regulator of reactive oxygen species (NRROS) as a novel interactor of Rubicon. Consistent with this interaction, *Rubcn*-mutant mice displayed markedly reduced oxidative damage, indicated by decreased lipid peroxidation after injury. Together, these findings indicate that Rubicon promotes acute neuroinflammatory and oxidative stress responses following TBI by modulating autophagy and ROS production. Rubicon mediated pathways may serve as therapeutic targets that offer a neuroprotective strategy to improve outcomes after TBI.

## Introduction

Traumatic brain injury (TBI) has one of the highest global incidences among neurological disorders and remains a leading cause of injury-related death and disability. (1) In addition to neuronal cell death and damage, TBI triggers pronounced neuroinflammation. Following the initial insult, resident microglia and infiltrating peripheral immune cells, including macrophages and monocytes, are recruited to the site of damage to drive inflammatory responses. (2) Although the initial neuroinflammatory responses are aimed at repairing the damage, they can exacerbate injury if not regulated properly.

Macro-autophagy (autophagy), a lysosome-dependent cellular catabolic pathway, plays a crucial role in the control of neurodegeneration and neuroinflammation after TBI. We previously demonstrated that autophagy is impaired in neurons and microglia during the acute phase of TBI. In neurons, this impairment induces ER stress and contributes to neuronal cell death. (3, 4) The pool of damage-associated molecular patterns (DAMPs) increases with more cell death, creating a highly pro-inflammatory milieu in the peri-injury region. (3) Autophagy also directly regulates innate immunity including the cGAS/STING pathway for the expression of pro-inflammatory type 1 interferon (IFN) genes, which sustain microglial activation, exacerbate neuronal injury, and promote chronic neurodegeneration in experimental mouse models of TBI. (5–7) Microglial type 1 IFN signaling has been in particular identified as a critical pathogenic driver of cortical neuroinflammation after TBI. (8) Therefore, autophagy is an essential pathway for regulation of inflammatory responses after TBI and understanding its regulation in this context may lead to development of novel targeted therapies.

A central coordinator of autophagy is Beclin 1 (BECN1), a component of the type III PI3 kinase necessary for the initiation of autophagosome biogenesis. Its binding partner, Rubicon (RUBCN), negatively regulates canonical autophagy by inhibiting autophagosome initiation, maturation, and lysosomal fusion.(9–14) Decline in autophagic activity with age is correlated with elevated Rubicon expression, which is implicated in age-related polyglutamine (polyQ) aggregation, interstitial fibrosis, and α-synuclein pathology. (15, 16) However, in addition to its role as a negative regulator of autophagy, Rubicon is also a positive regulator of conjugation of ATG/LC3 to single membranes (CASM) pathways, which regulate phagocytic and endosomal functions. (17–19) For instance, Rubicon is involved in the clearance of axonal and microglial debris through LC3-associated phagocytosis (LAP). (20, 21) Failure of LAP is reported to cause accumulation of myelin debris and apoptotic cells and failure of macrophage scavenger receptor 1 (MSR1) recycling in a murine model of multiple sclerosis. (22) Similarly, Rubicon-dependent LC3-associated endocytosis (LANDO) has been shown to participate in recycling of amyloid beta (Aβ) receptors in an Alzheimer’s disease model, leading to exacerbated amyloid deposition, plaque formation, and reactive microgliosis. (23, 24) The CASM pathways also regulate inflammatory signaling but the proposed roles appear context dependent. While LAP-deficient tumor associated macrophages upregulate type 1 IFN signaling via the STING pathway to promote anti-tumor responses, other murine models have shown that Rubicon promotes lupus autoimmunity. (25, 26) Together, these contrasting findings suggest that Rubicon exerts highly context-dependent effects on inflammation, leading to uncertainty regarding its role in inflammatory regulation.

Regulation of inflammation by Rubicon-dependent pathways is fundamentally linked to the production of reactive oxygen species (ROS). A major source of ROS is the nicotinamide adenine dinucleotide phosphate (NADPH) oxidase complex, NOX2. It is involved in regulation of phagocytosis and together with Rubicon participates in LAP. (27–29) While transient ROS generation helps recruit microglia to the site of injury for repair, excessive ROS production promotes oxidative stress due to depletion of endogenous antioxidants. (30) This can damage biomolecules through the peroxidation of membrane lipids and the oxidation of proteins and DNA. In myeloid cells, NOX2-mediated ROS is constrained by the negative regulator of reactive oxygen species (NRROS). NRROS limits ROS generation by facilitating degradation of Gp91*^phox^*, the catalytic subunit of NOX2, and suppression of phosphorylation of p47*^phox^*, the regulatory subunit. (31–33) Beyond ROS regulation, NRROS also contributes to the activation of TGFβ1 signaling, desensitization of TLR signaling, and development and maintenance of microglia. (34–37)

Given Rubicon’s highly context-dependent influence on inflammatory responses in different experimental paradigms, we investigated its regulatory role in TBI. Using a controlled cortical impact (CCI) model of TBI in *Rubcn*-mutant mice, we demonstrate that mutation in Rubicon markedly dampens acute neuroinflammatory responses. This transient suppression of inflammation was associated with improved motor recovery despite comparable cortical damage. We found that *Rubcn*-mutant mice exhibit attenuated injury-induced autophagy inhibition and reduced accumulation of damage-associated markers. Furthermore, proteomic analysis revealed that the mutant mice express a truncated Rubicon isoform lacking the RUN domain. Mechanistically, we identify NRROS as a novel Rubicon-interacting protein, which aligns with the diminished oxidative damage observed in *Rubcn*-mutant mice after injury. These findings suggest that Rubicon promotes acute neuroinflammation after TBI through coordinated downregulation of autophagy and exacerbation of oxidative stress.

## Results

### *Rubcn*-mutant mice have attenuated inflammatory gene expression following TBI

To examine whether Rubicon plays a role in neuroinflammation following TBI, we induced moderate brain injury using controlled cortical impact (CCI) model in Rubicon (*Rubcn*)-mutant and wild-type control mice. We collected RNA from peri-lesion ipsilateral cortices at 3 days post injury (dpi), the peak of both inflammatory responses and inhibition of autophagy in microglia and macrophages after TBI, for bulk RNA-sequencing. (3) Variant sparse Partial Least Squares-Discrimination Analysis (sPLS-DA) analysis demonstrated that the gene expression profiles of samples, including wild-type sham, wild-type TBI, *Rubcn*-mutant sham, and *Rubcn*-mutant TBI, clustered based on injury (55.2%, x-axis) and genotype (2.9%, y-axis) (**Fig. 1A**). Pair-wise comparisons between injured and sham groups within each genotype (**Fig. 1B, Tables S1-6**) identified many differentially expressed genes (DEGs).

**Figure 1:**
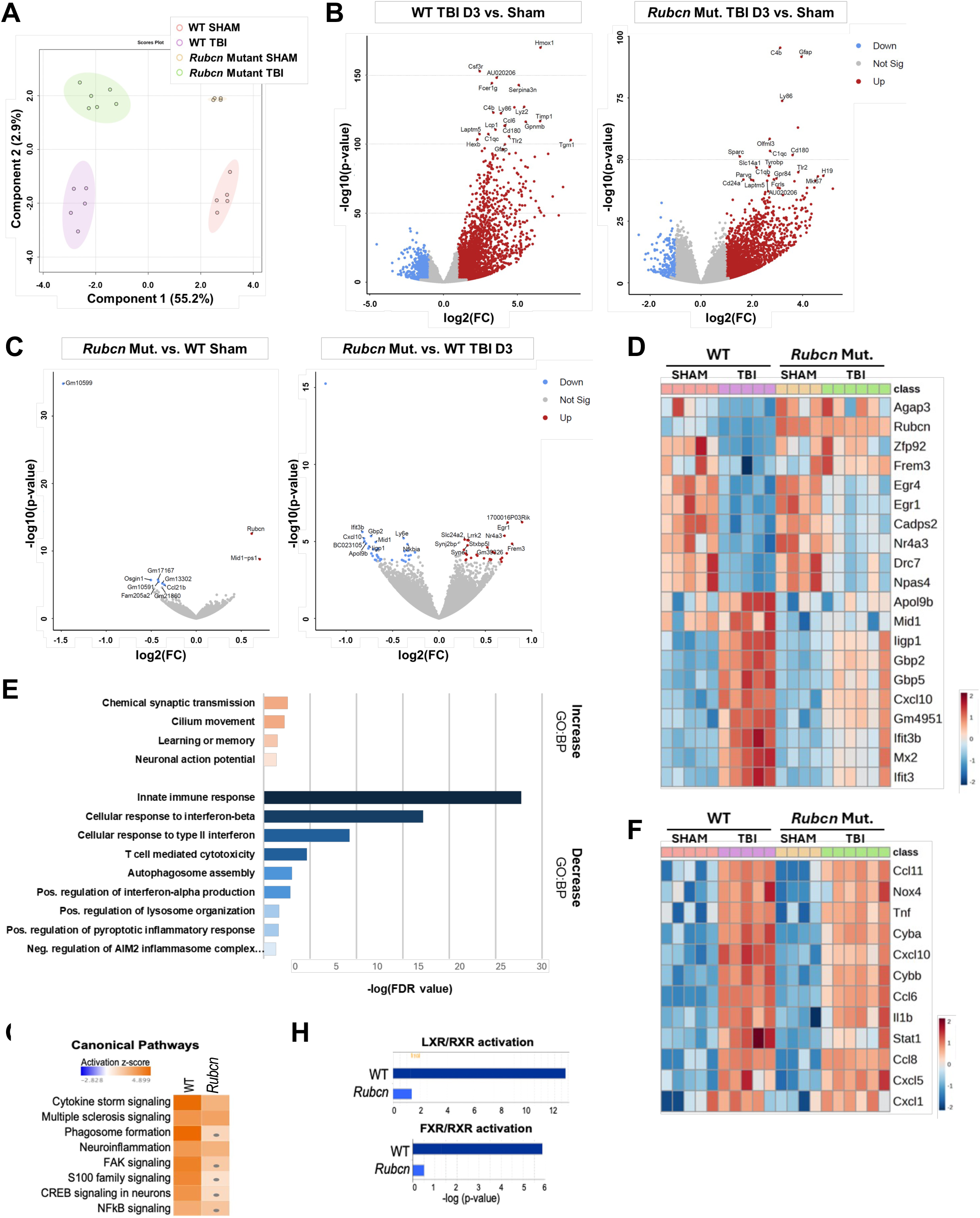
Rubcn-mutant mice have attenuated inflammatory gene expression following TBI. **A** Variant sparse Partial Least Squares-Discriminate Analysis (sPLS-DA) plot demonstrating group separation among ipsilateral cortical samples from wild-type sham, Rubcn-mutant sham, wild-type injured, and Rubcn-mutant injured mouse groups. **B** Volcano plots highlighting differentially expressed genes (DEGs) between injured and sham mice in wild-type (left) and Rubcn-mutant (right) samples, with red representing significantly upregulated genes and blue representing significantly downregulated genes. **C** Volcano plots showing DEGs between Rubcn-mutant and wild-type mice in sham (left) and injured samples (right). A data point corresponding to the Rubcn gene has been removed from the right volcano plot to minimize the y-axis. **D** Heatmap of gene expression across the samples demonstrating top upregulated (top-half) and downregulated (bottom-half) DEGs between Rubcn-mutant and wild-type TBI mice. **E** Gene Ontology Biological Process (GO:BP) analysis of significant DEGs (P-value<0.01) between Rubcn-mutant and wild-type TBI samples. Top graph shows GO-terms associated with upregulated genes (in orange) and bottom graph shows GO-terms associated with downregulated genes (in blue) in Rubcn-mutant over wild-type TBI samples with FDR<0.05. **F** Heat map of selected TBI-induced inflammatory DEG expression across wild-type and Rubcn-mutant sham and TBI samples. **G** Ingenuity Pathway Analysis (IPA) analysis comparing canonical pathways activated following TBI in wild-type versus Rubcn-mutant mice. Orange (activated) and blue (inhibited) indicate significantly enriched canonical pathways based on activation z-score, while gray dots represent pathways detected in the specified dataset that did not meet significance thresholds. **H** IPA analysis of lipid-related pathways activated following TBI in wild-type versus Rubcn-mutant mice. Sample size (n) were as follows: WT Sham=5, WT TBI D3=5, Rubcn Mut. Sham=4, Rubcn Mut. TBI D3=6.

However, the significance of upregulated DEGs in *Rubcn*-mutant injured mice was generally lower than in wild-type injured mice. While pair-wise comparisons between *Rubcn*-mutant and wild-type mice within sham and injured groups identified overall fewer DEGs, the genotype effect was stronger under injury conditions (**Fig. 1C**). This was also reflected in comparison of top upregulated and downregulated DEGs between injured mice of both genotypes, which confirmed that injury effect was attenuated in *Rubcn*-mutant mice (**Fig. 1D**). Further analyses of DEGs with Gene Ontology Biological Process (GO: BP) revealed that the GO-terms associated with genes expressed at lower levels in injured *Rubcn*-mutant mice were enriched for immune responses and immune regulation (**Fig. 1E, Tables S7-8**) suggesting attenuated inflammation following injury. These responses could be attributed to the reduced induction of TBI-associated inflammatory DEGs in the mutant mice (**Fig. 1F**). When comparing the effect of injury over sham within each genotype, Ingenuity Pathway Analysis indicated that while activation of canonical inflammatory signaling pathways was observed in both groups, it was markedly stronger in the wild-type as compared to *Rubcn*-mutant injured mice (**Fig. 1G, Tables S9-12**). Similarly, lipid metabolism-related pathways were less robustly induced in the *Rubcn*-mutant mice after injury (**Fig. 1H**).

Microglia and macrophages play an important role in TBI neuroinflammation. Analysis of our RNA-seq data indicated overall decrease in markers associated with microglial and macrophage activation in *Rubcn*-mutant as compared to wild-type TBI mice (**Figure S1A-B**). The decrease was more pronounced for pro-inflammatory (M1-like) genes as compared to anti-inflammatory (M2-like) genes. Attenuated expression of ionized calcium-binding adaptor molecule 1 (IBA1), a marker of both microglia and infiltrating monocytes, was observed in the ipsilateral cortex of *Rubcn*-mutant mice at 3 dpi (**Fig. S1C-D**). No changes were observed for astrocyte marker glial fibrillary acidic protein (GFAP).

**Supplemental Figure 1:**
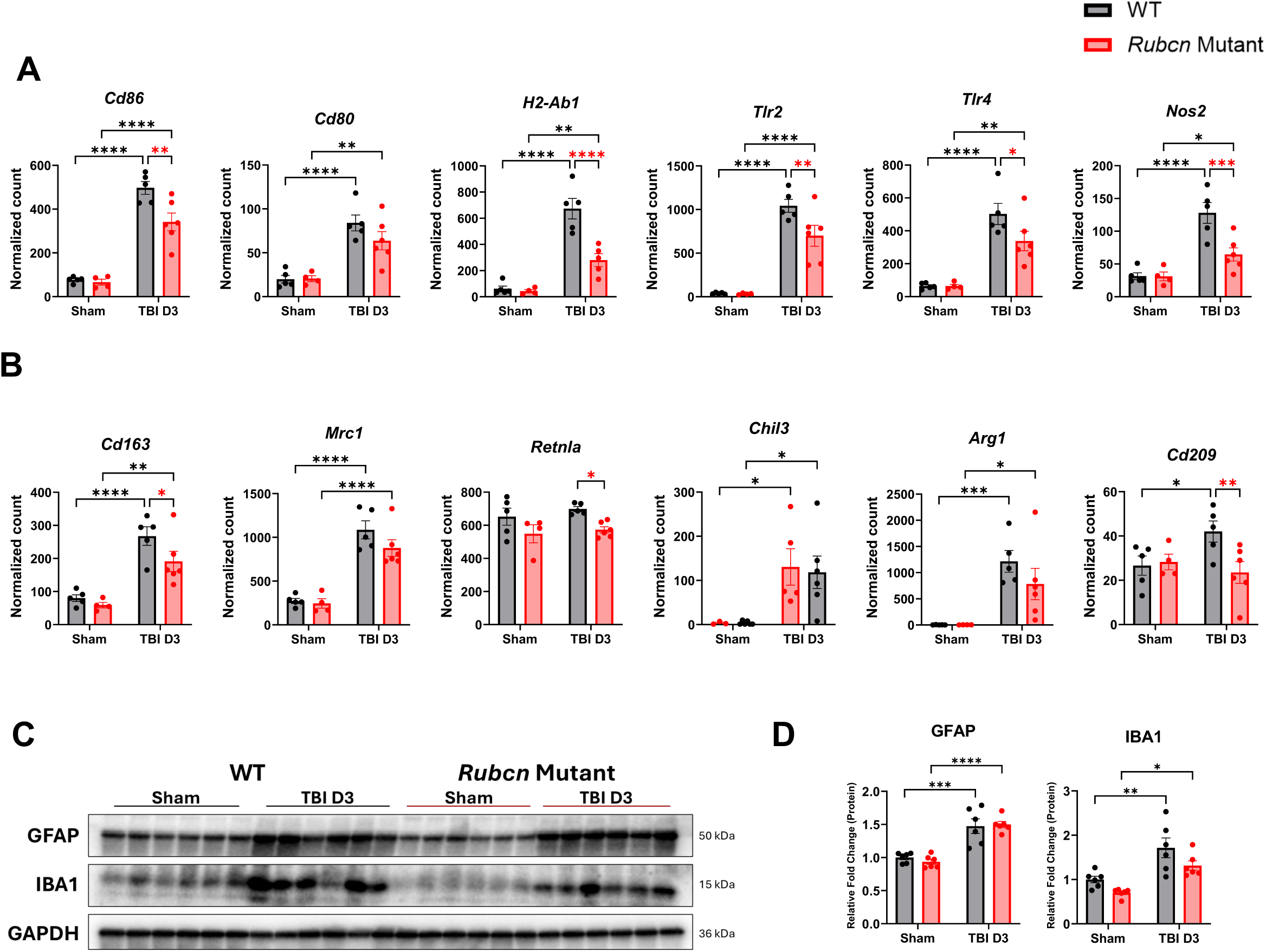
Rubicon mutation decreases microglial pro-inflammatory polarization. **A** Normalized count of pro-inflammatory (M1-like) mRNA in wild-type and *Rubcn*-mutant mice after day 3 (TBI D3) of injury in comparison to sham. **B** Normalized count of anti-inflammatory (M2-like) mRNA in wild-type and *Rubcn*-mutant mice after day 3 (TBI D3) of injury in comparison to sham. **C** Immunoblot comparing ipsilateral cortices from wild-type and *Rubcn*-mutant mice after day 3 of injury to sham. **B** Densitometric quantification of immunoblot in **D**. Bars represent mean±SEM. Sample size (n) for **A-B** were as follows: WT Sham = 5, WT TBI D3=5, *Rubcn* Mut. Sham=4, *Rubcn* Mut. TBI D3=6. Statistical analyses: Two-way ANOVA with post-hoc Uncorrected Fisher’s LSD test. Sample size (n) for **C-D** were as follows: WT Sham = 6, WT TBI D3=6, *Rubcn* Mut. Sham=6, *Rubcn* Mut. TBI D3=6. Statistical analyses: Two-way ANOVA with post-hoc Tukey’s test. Significance is represented by * p-value<0.05, ** p-value<0.005, *** p-value<0.0005, **** p-value<0.0001.

### Reduction in TBI-induced inflammation in *Rubcn*-mutant mice is temporary

We used a separate set of samples to confirm RNA-seq data and determine the time course of these changes. Our real-time qPCR data confirmed attenuated inflammatory and lipid-metabolism related responses in *Rubcn*-mutant mice after injury at 3 dpi (**Fig 2A-B**). However, with the exception of *Pparg,* no significant differences were apparent at 1 dpi. Neuroinflammation and microglial activation can persist long-term after injury. (38) At 29 dpi, corresponding to the time of transition from sub-acute to chronic TBI, both wild-type and Rubicon mutant mice continued to exhibit elevated expression of inflammatory genes (**Fig 2C-D**). However, the differences between the two genotypes had largely diminished by this time point. Overall, our data indicate that Rubicon is involved in regulation of acute but not long-term inflammation following TBI.

**Figure 2:**
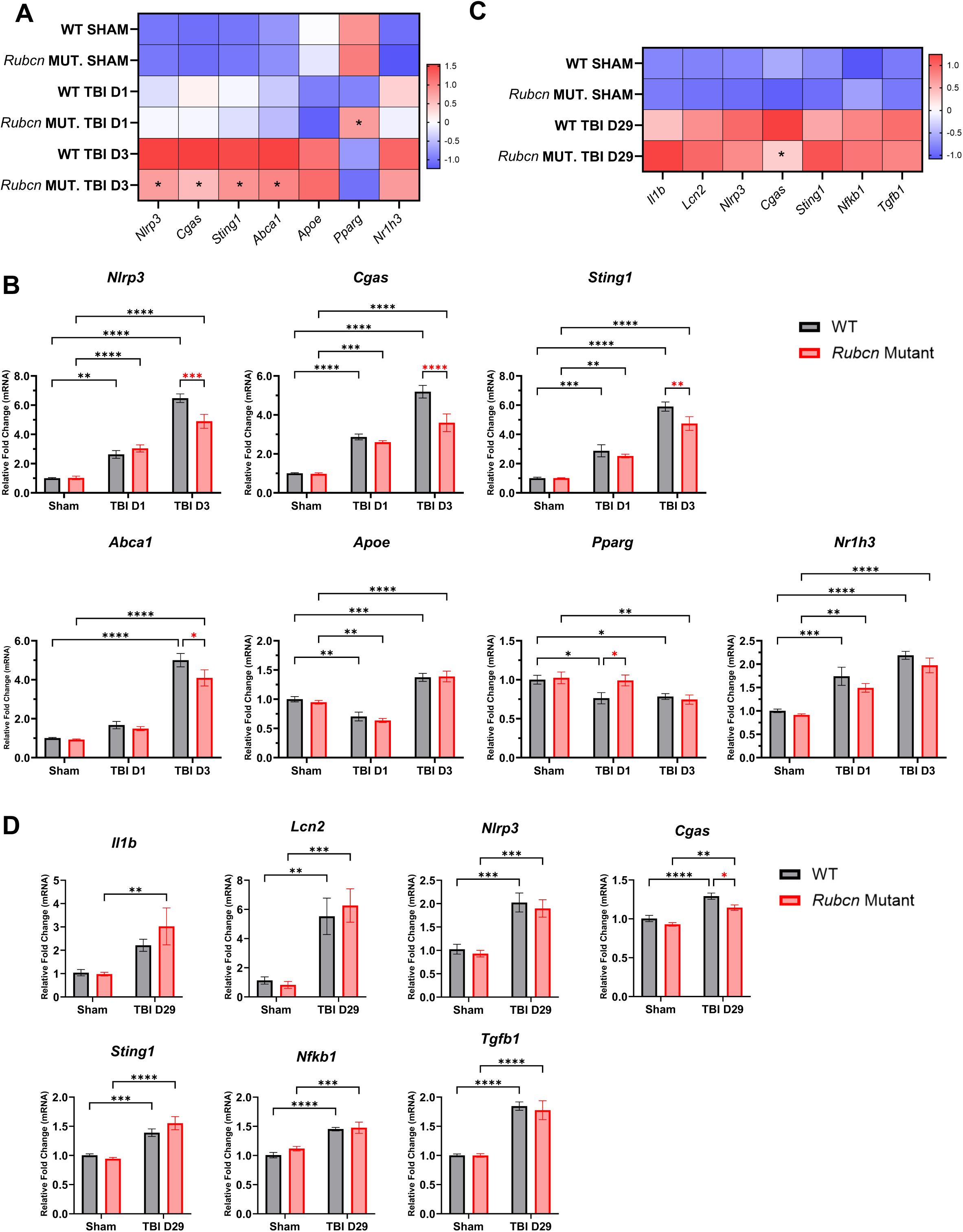
Reduction in TBI-induced inflammation in Rubcn-mutant mice is temporary. **A** Heatmap of RT-qPCR data showing expression of inflammatory genes in ipsilateral cortices of wild-type and *Rubcn*-mutant mice at 1 dpi (TBI D1) and 3 dpi (TBI D3) in comparison to sham. Color coding is based on z-score. **B** Fold change in expression levels of indicated genes in wild-type and *Rubcn*-mutant mice corresponding to **A**. **C** Heatmap of RT-qPCR data showing expression of inflammatory genes in ipsilateral cortices of wild-type and *Rubcn*-mutant mice at 29 dpi (TBI D29). **D** Fold change in expression levels of indicated genes in wild-type and *Rubcn*-mutant mice corresponding to C. Bars represent mean ± s.e.m. Sample size (*n*) for A-B were as follows: WT Sham=8, WT TBI D1=5, WT TBI D3=9, *Rubcn* Mut. Sham=8, *Rubcn* Mut. D1=7, *Rubcn* Mut. D3=9. Sample size (*n*) for C-D were as follows: WT Sham=6, WT TBI D29=6, *Rubcn* Mut. Sham=7, *Rubcn* Mut. TBI D29=5. Statistical analyses for all data: Two-way ANOVA with post hoc Tukey’s test and significance is represented by * *P*-value<0.05, ** *P*-value <0.005, *** *P*-value <0.0005, **** *P*-value <0.0001.

### Rubicon mutant mice show improved motor recovery

To evaluate whether Rubicon influences functional recovery after injury, we monitored recovery of motor coordination. *Rubcn*-mutant mice performed better on the beam walk test, exhibiting fewer foot faults overall, with significant reduction at 3 and 14 dpi (**Fig. 3A**). Gait analysis using CatWalk demonstrated that *Rubcn*-mutant mice displayed increased one (running) to two (normal walk) limb support and decreased three to four limb support, indicative of improved stability and coordination (**Fig. 3B**). Cortical damage was assessed by quantifying the percent of ipsilateral cortex remaining at 29 dpi. No significant differences were observed between wild-type and *Rubcn*-mutant mice (**Fig. 3C**). These findings suggest 8 that while decreased inflammatory responses in *Rubcn*-mutant mice are temporary, they result in enhanced motor recovery after brain injury.

**Figure 3:**
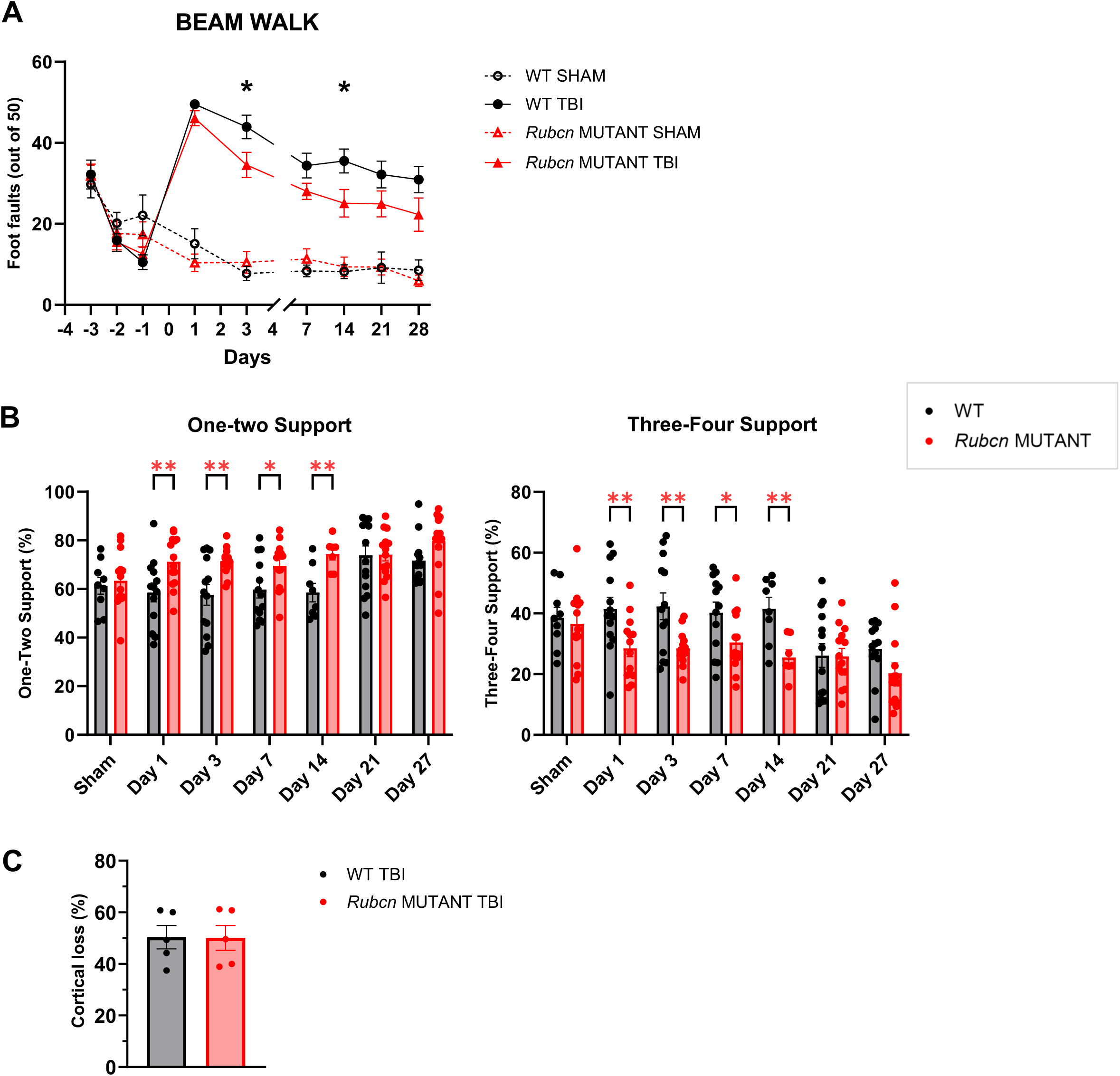
Rubicon mutant mice show improved motor recovery. **A** Number of foot faults before and after injury in wild-type and Rubcn-mutant mice by beam walk test. **B** Percent of one/two-foot support (left) and three/four-foot support (right) after injury in wild-type and Rubcn-mutant mice by assessed by CatWalk test. **C** Percentage of ipsilateral cortical damage at 29 dpi between injured wild-type and Rubcn-mutant mice. Data represent mean ± s.e.m. Sample size (n) for A were as follows: WT Sham=11, WT TBI=13, Rubcn Mut. Sham=13, Rubcn Mut. TBI=14. Sample size (n) for B were as follows: n=7-14 mice per group as indicated. Sample size (n) for C were as follows: WT TBI=5, Rubcn Mut. TBI=5. Statistical Analyses: For A-B, mixed-effects analysis with post hoc Tukey’s test; For C, Unpaired Student’s t-test. For all data, significance is represented by * P-value<0.05, ** P-value<0.005.

### Rubicon mutation decreases injury-induced autophagy inhibition

We investigated the mechanisms of how mutation in Rubicon may lead to decrease in inflammation and improved TBI recovery. Rubicon is a negative regulator of autophagy, and decreased Rubicon levels are associated with increased autophagic activity. (15, 16) We previously demonstrated that during the acute phase after TBI, autophagy is inhibited in microglia and macrophages contributing to increased inflammation. (3, 4) To determine whether Rubicon regulates injury-induced inflammation through its effects on autophagy, we examined autophagy and damage-associated markers in the ipsilateral cortex and hippocampus at 1 and 3 dpi. We observed robust accumulation of P62/SQSTM1 protein indicative of autophagy inhibition in the peri-lesion cortex of wild-type mice starting at 1 dpi (**Fig. S2A-B**), which persisted through 3 dpi (**Fig. 4A-B**). In contrast, *Rubcn*-mutant mice showed minimal P62/SQSTM1 accumulation at either time point. At the transcriptional level, *Sqstm1* gene expression was slightly upregulated in wild-type mice at 1 dpi (**Fig. S2C**). This was, however, insufficient to account for the pronounced accumulation of P62/SQSTM1 protein, indicating protein-level regulation as we have reported previously. (4) Levels of autophagosome associated LC3-II protein were similarly higher in injured wild-type mice at 3 dpi but unaltered in injured *Rubcn* mutants (**Fig.4A-B, S2A-B**). No changes were observed in *Map1lc3b* gene expression (**Fig. S2C**). We also assessed damage-associated indicators and found that α-fodrin protein levels were reduced in *Rubcn*-mutant mice at 3 dpi (**Fig.4A-B, S2A-B).**

**Figure 4:**
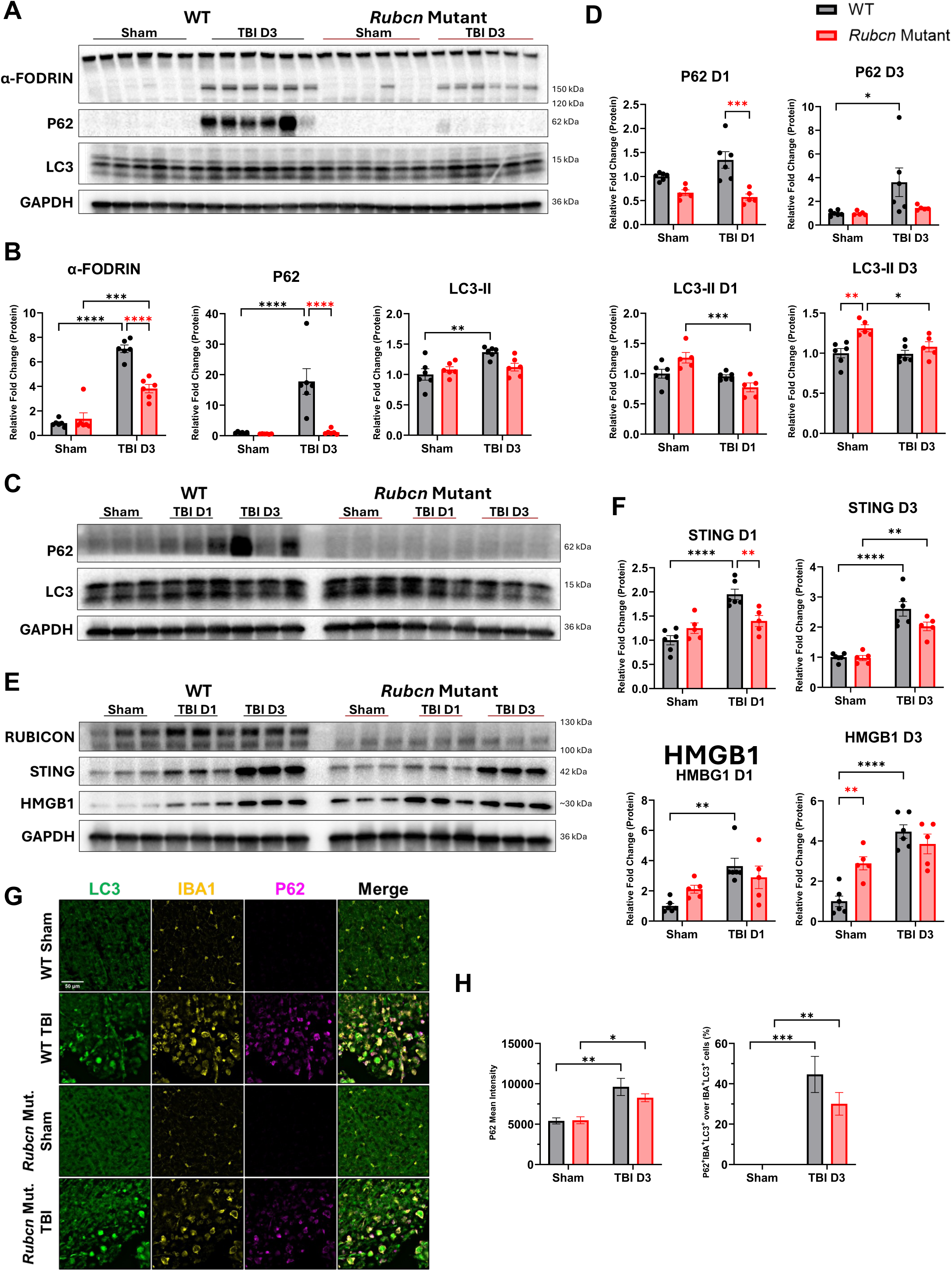
Rubicon mutation decreases injury-induced autophagy inhibition. **A** Immunoblot comparing damage-associated marker (α-fodrin) and autophagy markers (P62, LC3II) in ipsilateral cortices from wild-type and Rubcn-mutant mice at 3 dpi to sham. **B** Densitometric quantification of immunoblot in A. **C** Immunoblot of ipsilateral hippocampus at 1 and 3 dpi in wild-type and Rubcn-mutant mice probed with autophagy markers. **D** Densitometric quantification of immunoblot in C. **E** Immunoblot of ipsilateral hippocampus at 1 dpi (TBI D1) and 3 dpi (TBI D3) in wild-type and Rubcn-mutant mice probed with damage-associated markers. **F** Densitometric quantification of immunoblot in E. **G** Immunofluorescence (IF) images of GFP-LC3 (green) coronal sections from wild-type and Rubcn-mutant sham and injured mice at 3 dpi stained for IBA1 (in yellow) and P62 (in magenta). Scale bar represents 50 μm. **H** Quantification of mean fluorescence intensity of P62 (left) and percent of P62/SQSTM1+ cells within LC3+/IBA+ cells (right) from G. All bars represent mean ± s.e.m. Sample size (n) for A-B were as follows: WT Sham=6, WT TBI D3=6, Rubcn Mut. Sham=6, Rubcn Mut. D3=6. Sample size (n) for C-F were as follows: WT Sham=6, WT TBI D1=6, WT TBI D3=6, Rubcn Mut. Sham=5, Rubcn Mut. D1=5, Rubcn Mut. D3=5. Sample size (n) for G-H were as follows: WT Sham=4, WT TBI D3=4, Rubcn Mut. Sham=4, Rubcn Mut. D3=4. Statistical analyses for all data: Two-way ANOVA with post hoc Tukey’s test and significance is represented by * P-value<0.05, ** P-value<0.005, *** P-value<0.0005, **** P-value<0.0001.

**Supplemental Figure 2:**
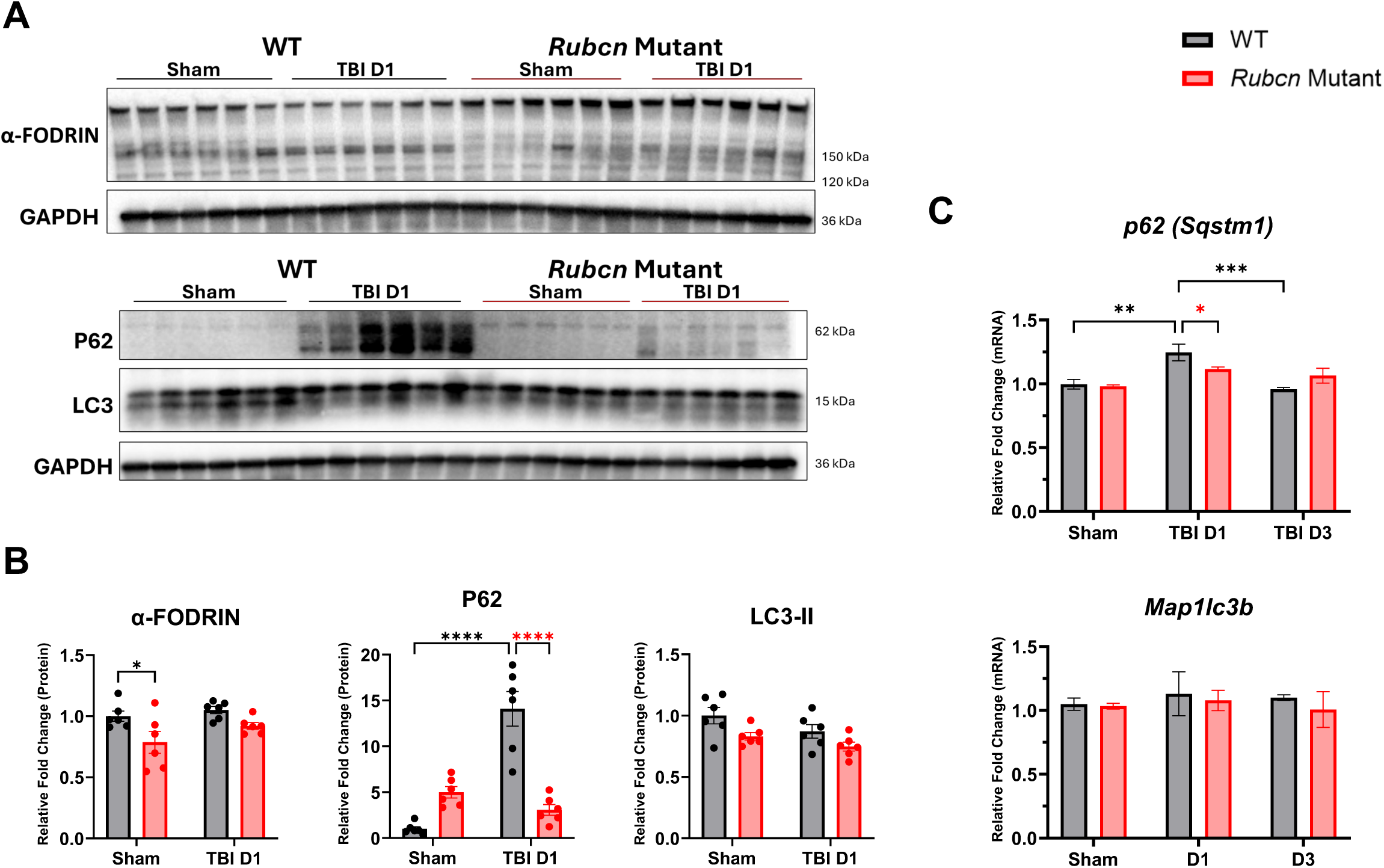
Rubicon mutation decreases injury-induced autophagy inhibition. **A** Immunoblot comparing damage-associated marker (top) and autophagy markers (bottom) in ipsilateral cortices from wild-type and *Rubcn*-mutant mice after 1 day post injury (dpi).**B** Densitometric quantification of immunoblot in **A**. **C** Fold change of mRNA levels of *Sqstm1* and *Map1lc3b* in wild-type and *Rubcn*-mutant mice after 1 and 3 dpi compared to sham. All bars represent mean±SEM. Sample size (n) for **A-B** were as follows: WT Sham = 6, WT TBI D3=6, *Rubcn* Mut. Sham=6, *Rubcn* Mut. D3=6. Sample size (n) for **C** were as follows: WT Sham = 3, WT TBI D3=3, *Rubcn* Mut. Sham=3, *Rubcn* Mut. D3=3. Statistical analyses: Two-way ANOVA with post-hoc Tukey’s test and significance represented by * p-value<0.05, ** p-value<0.005, *** p-value<0.0005, **** p-value<0.0001.

A similar genotype-dependent pattern of reduced P62/SQSTM1 accumulation was detected in the ipsilateral hippocampus (**Fig. 4C-D**), showing significance at 1 dpi. Wild-type mice also exhibited significant accumulation of HMGB1, another damage-associated marker, in the hippocampus at both 1 and 3 dpi (**Fig. 4E-F**), while *Rubcn*-mutant mice showed no injury-induced increase. Consistent with decreased inflammation, STING protein levels increased in the hippocampal tissue of wild-type but not *Rubcn*-mutant mice at 1 dpi **(Fig. 4E-F**). These results suggest that *Rubcn*-mutant mice exhibit reduced autophagy inhibition and attenuated tissue damage following injury.

We previously demonstrated that autophagy is specifically inhibited in microglia and macrophages after TBI. (3) To assess microglia/macrophage specific regulation of autophagy, we performed immunostaining for IBA1 to mark microglia and monocytes with P62/SQSTM1 on coronal brain sections from GFP-LC3 transgenic wild-type and *Rubcn*-mutant mice at 3 dpi. (39) As previously, we detected a significant increase in P62/SQSTM1 within IBA^+^ cells and an increased percentage of P62/SQSTM1^+^ cells within LC3^+^/IBA^+^ populations in both injured groups compared with sham controls (**Fig. 4G-H**).(3) However, *Rubcn*-mutant mice displayed less pronounced overall and IBA1^+^ cell-specific accumulation of P62/SQSTM1, indicating attenuated inhibition of microglial/monocyte autophagy. To further investigate the effect of Rubicon mutation on autophagy, we cultured primary bone marrow derived macrophages (BMDMs) from wild-type and *Rubcn*-mutant mice and measured autophagy flux under baseline and starvation conditions. (40) *Rubcn*-mutant BMDMs displayed elevated baseline autophagy but no difference in starvation-induced autophagy relative to wild-type controls (**Fig. S3A-B**), suggesting context-specific regulation of autophagy by Rubicon.

### Truncated RUBCN protein is present in *Rubcn*-mutant mice

Full length RUBCN protein is expected to migrate at approximately 130 kDa. Previous studies reported a faster migrating RUBCN protein at 100 kDa which represented an alternative isoform translated through internal ribosome entry sites. (41, 42) This smaller isoform was present in cells from both wild-type and conditional *Rubcn*-mutant mice used in that study. Consistent with these reports, we detected RUBCN signal in wild-type mice at both ∼130 kDa and ∼100 kDa (**Fig. 5A, S4A-B**). In *Rubcn*-mutant mice, the 130 kDa band was absent as expected, whereas the 100 kDa band persisted. To determine whether this corresponded to the truncated RUBCN protein, we conducted immunoprecipitation (IP) of endogenous Rubicon from brain lysates of uninjured mice using two different antibodies, Rubicon A (RbA) and Rubicon B (RbB). The 100 kDa and 130 kDa RUBCN bands were detected with both RbA and RbB; however, a background signal at ∼150 kDa was also observed with RbB (**Fig. S4A-B**). RUBCN was successfully immunoprecipitated from both wild-type and *Rubcn*-mutant mice (**Fig. 5B**) with wild-type samples showing 130 kDa and 100 kDa and *Rubcn* mutant only the 100 kDa band. Neither band was present when IgG control antibody was used. All samples were subjected to mass spectrometry (MS). In wild-type samples, we detected peptides p1 to p11 spanning the entire length of RUBCN amino-acid sequence (**Fig. 5C, Tables S13-14**). Analysis of *Rubcn*-mutant samples revealed that while the mutant RUBCN protein lacked peptides corresponding from p1 to p5 at the N-terminus, it retained peptides beginning at p6 (**Fig. 5D, S4C**). *Rubcn* gene sequence analysis identified a potential alternative start site at Met^296^ between peptides 5 and 6 (**Fig. 5C**). This is consistent with prior reports of internal Rubicon translation initiation in cultured cells. (41, 42)

**Figure 5:**
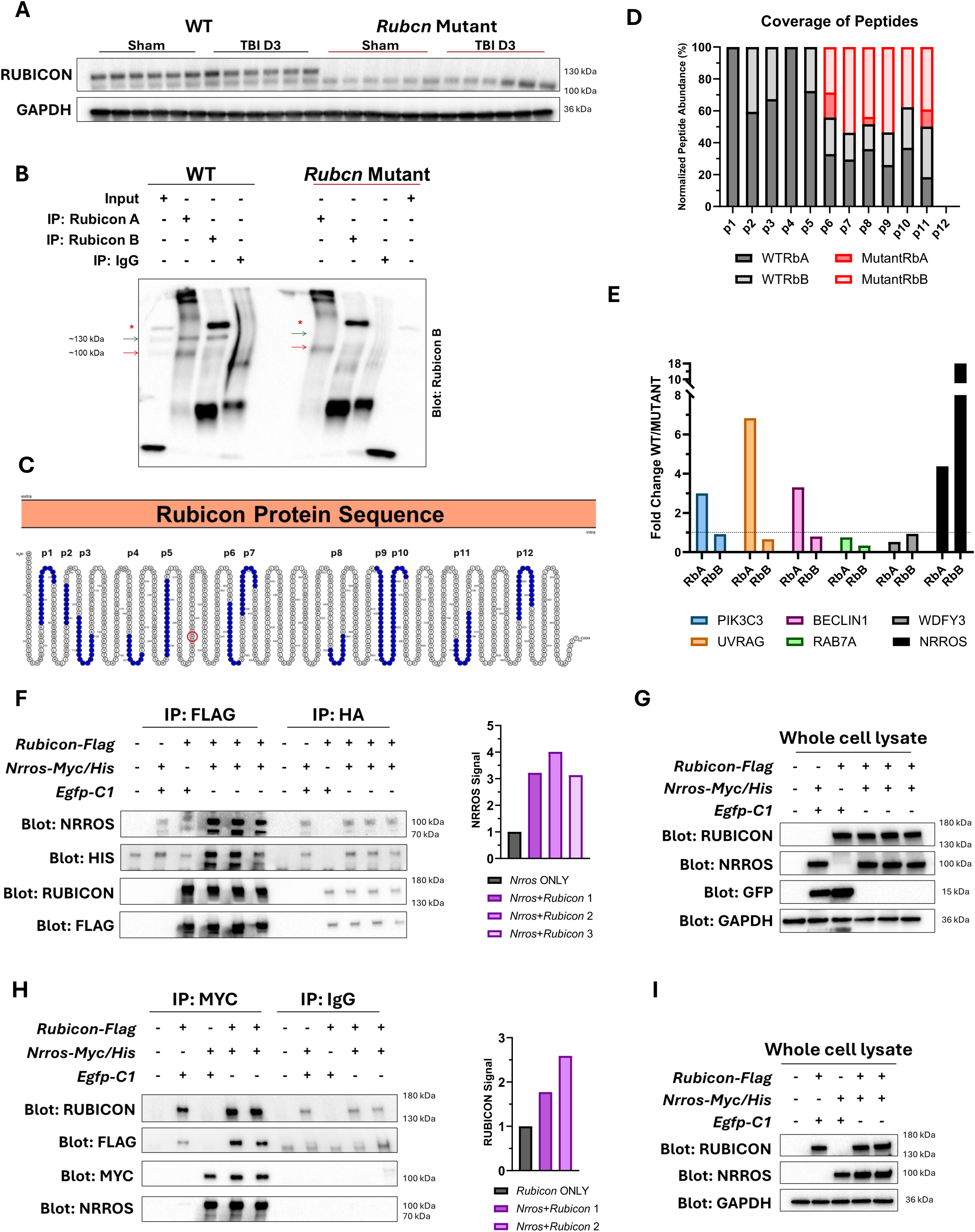
RUBCN protein interacts with NRROS *in vivo* and *in vitro*. **A** Immunoblot comparing lysates of ipsilateral cortices from wild-type and Rubcn-mutant mice probed with Rubicon A (RbA) antibody. **B** Western blot showing immunoprecipitation of Rubicon protein from brain lysates of wild-type and Rubcn-mutant mice using Rubicon A (RbA) and Rubicon B (RbA) antibodies. * represents background signal. Blot is probed with RbB antibody. **C** Rubicon protein (amino-acid) sequence highlighting peptides (p1-p12 in blue) used for peptide mapping of wild-type and mutant RUBCN proteins. Red circle is highlighting Met296 as a potential alternative translation initiation site. **D** Normalized peptide abundance across the length of RUBCN protein in wild-type and Rubcn-mutant immunoprecipitates using RbA and RbB antibodies. **E** Comparative enrichment analysis showing fold change of proteins enriched in wild-type over Rubcn-mutant immunoprecipitates using RbA and RbB antibodies. **F** Immunoprecipitation assay showing lysates of HEK293T cells transfected with indicated plasmids for 24 h and immunoprecipitated with anti-FLAG or anti-HA agarose beads. Left: Immunoblot of immunoprecipitates with the indicated antibodies. Right: Fold change of NRROS signal in EGFP-Rubicon-Flag and Nrros-Myc/His (Nrros+Rubicon1-3) co-transfected cells to EGFP-C1 and Nrros-Myc/His (Nrros ONLY) co-transfected cells. Each FLAG IP sample was normalized to corresponding HA IP sample for quantification. Sample size (n) = 3 biological replicates for EGFP-Rubicon-Flag and Nrros-Myc/His co-transfected cells. **G** Immunoblot of whole cell lysates used for immunoprecipitation assay in F. **H** Immunoprecipitation assay showing lysates of HEK293T cells transfected with indicated plasmids for 24 h and immunoprecipitated with anti-MYC or anti-IgG antibodies. Left: Immunoblot of immunoprecipitates with the indicated antibodies. Right: Fold change of RUBCN signal in EGFP-Rubicon-Flag and Nrros-Myc/His (Nrros+Rubicon1-2) co-transfected cells to EGFP-C1 and EGFP-Rubicon-Flag (Rubicon ONLY) co-transfected cells. Each MYC IP sample was normalized to corresponding IgG IP sample. Sample size (n) = 2 biological replicates for EGFP-Rubicon-Flag and Nrros-Myc/His co-transfected cells. **I** Immunoblot of whole cell lysates used for immunoprecipitation assay in H.

**Supplemental Figure 3:**
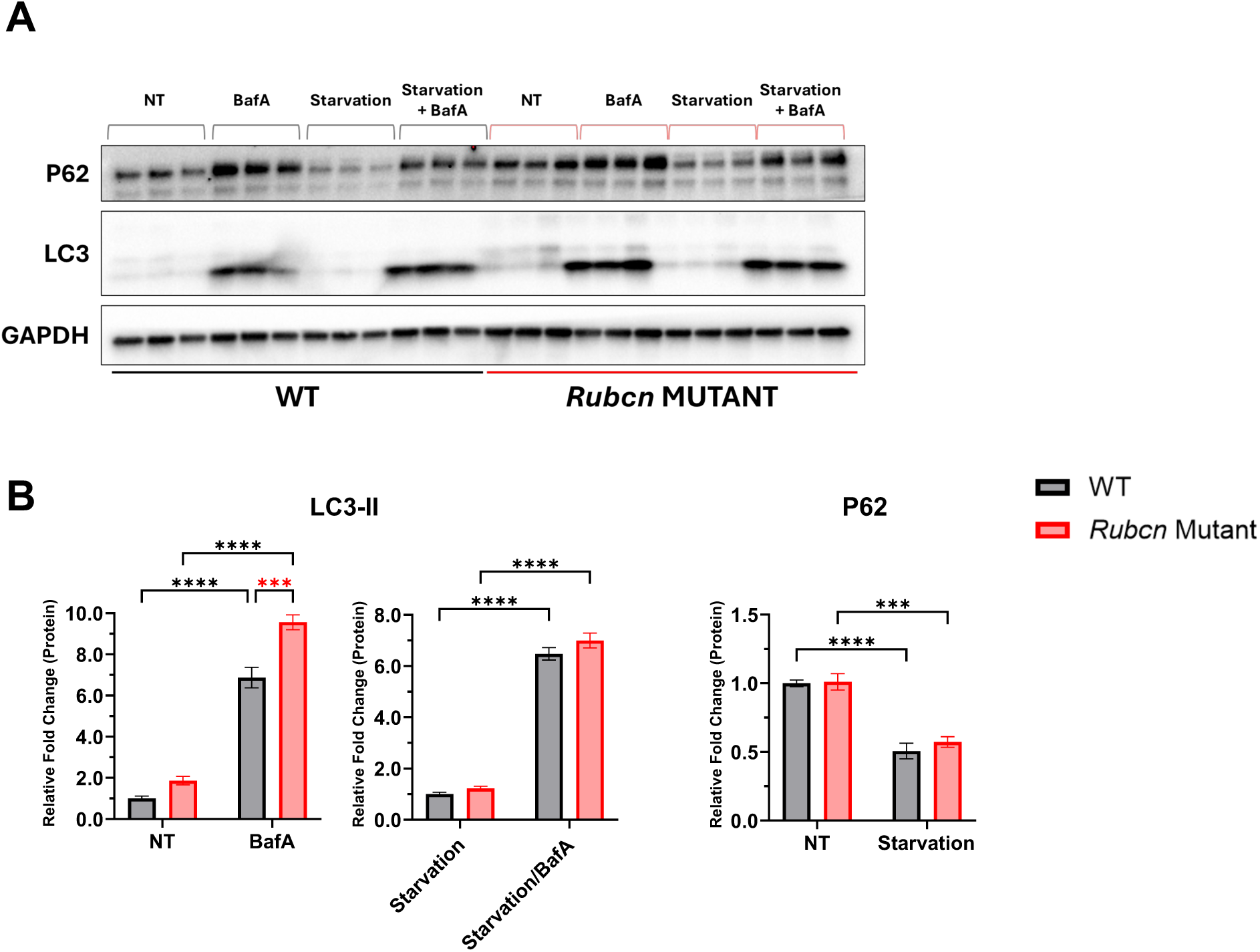
Rubicon mutation increases baseline autophagy flux *in vitro*. **A** Immunoblot of wild-type and *Rubcn*-mutant bone marrow derived macrophages (BMDMs) pre-treated with 50 nM Bafilomycin A (BafA) for an hr. and/or serum-starved (Starvation) for 4 hrs. NT represents no treatment group. **B** Densitometric analysis from **A.** Data represent mean±SEM. Sample size (n) were as follows: n=3 per group. Statistical analyses: Two-way ANOVA with post-hoc Uncorrected Fisher’s LSD test and significance represented by * p-value<0.05, ** p-value<0.005, *** p-value<0.0005, **** p-value<0.0001.

**Supplemental Figure 4:**
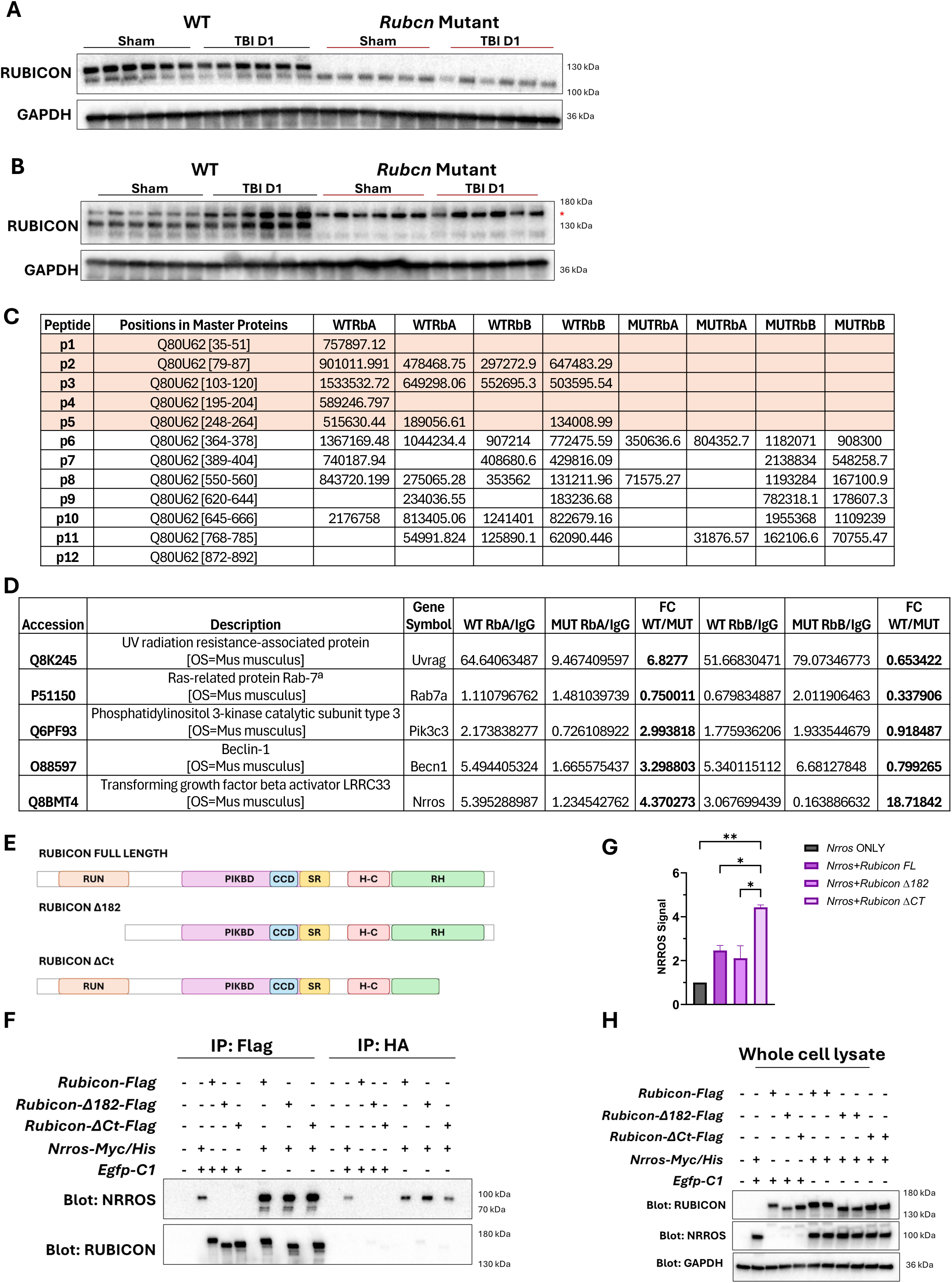
RUBCN protein interacts with NRROS *in vivo* and *in vitro*. **A** Immunoblot comparing lysates of ipsilateral cortices from wild-type and *Rubcn*-mutant mice probed with Rubicon A (RbA) antibody. **B** Immunoblot comparing lysates of ipsilateral cortices from wild-type and *Rubcn*-mutant mice probed with Rubicon A (RbB) antibody. * represents background signal. **C** Table showing normalized peptide abundance queried for peptides p1 to p12 across the length of RUBCN protein in wild-type and *Rubcn*-mutant immunoprecipitates using RbA and RbB antibodies. n=2 biological replicates for each genotype. **D** Analysis of comparative enrichment of proteins in wild-type over *Rubcn*-mutant immunoprecipitates using RbA and RbB antibodies. **E** Schematic diagram showing full-length and truncated mutants of RUCBN proteins and their protein domains. **F** Immunoprecipitation assay showing lysates of Hek293T cells transfected with indicated plasmids for 24 hrs. and immunoprecipitated with anti-FLAG or anti-HA agarose beads. Immunoprecipitates were subjected to western blot with the indicated antibodies. **G** Fold change of NRROS signal in *Nrros*+*Rubicon(FL, Δ182, or ΔCT*) transfected cells to *Nrros* ONLY transfected cells. Each FLAG IP sample was normalized to corresponding to HA IP sample. **H** Immunoblot of whole cell lysates used in immunoprecipitation assay in **F**. Data represent mean±SEM. Sample size (n) were as follows: n=2 biological replicates. Statistical analyses: One-way ANOVA with post-hoc Tukey’s test and significance represented by * p-value<0.05, ** p-value<0.005.

### RUBCN interacts with negative regulator of ROS (NRROS) protein

To identify novel RUBCN interactors that may mediate its eFect on TBI neuroinflammation, we analyzed our MS data for proteins enriched in wild-type but not *Rubcn*-mutant IP samples. Known interactors of RUBCN protein, including phosphatidylinositol 3-kinase catalytic subunit type 3 (PIK3C3 or VPS34), UV radiation resistance-associated gene (UVRAG), and BECN1, were preferentially enriched in wild-type samples (**Fig.5E, S4D, Table S15**), serving as internal quality controls for immunoprecipitation of full-length RUBCN. These findings are consistent with domain-specific interactions: PIK3C3 interacts with the RUN domain at the N-terminus of RUBCN (absent in *Rubcn*-mutant), UVRAG and BECN1 bind the central PIKBD domain (truncated). (43) In contrast, enrichment of Ras-related protein Rab-7a (RAB7A), another known RUBCN interactor, and autophagy-related WD repeat and FYVE domain-containing protein 3 (WDFY3) was weak and comparable between genotypes (**Fig.5E, S4D**). Our analysis also identified negative regulator of reactive oxygen species (NRROS) as a protein that was strongly enriched in wild-type but reduced in mutant Rubicon samples (**Fig.5E**).

To validate the association of NRROS with RUBCN, we overexpressed EGFP/Flag-tagged full-length RUBCN and Myc/His-tagged NRROS in mammalian cells to assess their interaction using immunoprecipitation (IP). (12, 44) We observed robust co-IP of NRROS with full-length RUBCN (**Fig. 5F-G**), and conversely, full-length RUBCN was recovered with NRROS (**Fig.5H-I**), confirming interaction between the two proteins. Both direct and indirect NRROS immunoprecipitates displayed an additional 70 kDa band below the expected 100 kDa band (**Fig. 5F & H**), which may represent a post-translationally modified form of NRROS, although this will need to be confirmed by additional studies. (45) The 70 kDa band was minimal or absent in whole cell lysates (**Fig. 5G & I**).

To further delineate the domain of Rubicon responsible for NRROS binding, we tested truncated Rubicon variants, RUBCNΔ182 (lacks the N-terminal RUN domain) and RUBCNΔCt (has truncated C-terminal RH domain, **Fig. S4E**).(46) Both mutants retained the ability to interact both the 70 kDa and 100 kDa NRROS species (**Fig. S4F-G**). These results suggest that NRROS interaction site lies between the end of the RUN domain and mid-region of RH domain, in the center of the Rubicon protein. Further domain mapping using additional truncation constructs will be required to precisely define the interaction interface.

### Rubicon mutation attenuates oxidative damage after TBI

Given the role of NRROS in the regulation of ROS and its preferential association with RUBCN in wild-type as compared to mutant mice, we hypothesized that the level of ROS production and consequent oxidative damage may diFer between the two genotypes following injury. (31) To assess oxidative damage, we measured levels of the lipid peroxidation marker, 4-hydroxynonenal (4-HNE). Previous studies have shown that 4-HNE levels peak between 48 and 72 hours post injury.(47) Consistent with this, wild-type mice exhibited significantly elevated 4-HNE levels at both 1 and 3 dpi (**Fig.6A-D**). *Rubcn*-mutant mice displayed reduced 4-HNE accumulation at both time points. Decreased oxidative damage in *Rubcn*-mutant TBI mice suggests that regulation of ROS may be a key mechanism in mediating Rubicon-dependent inflammatory responses following injury.

**Figure 6:**
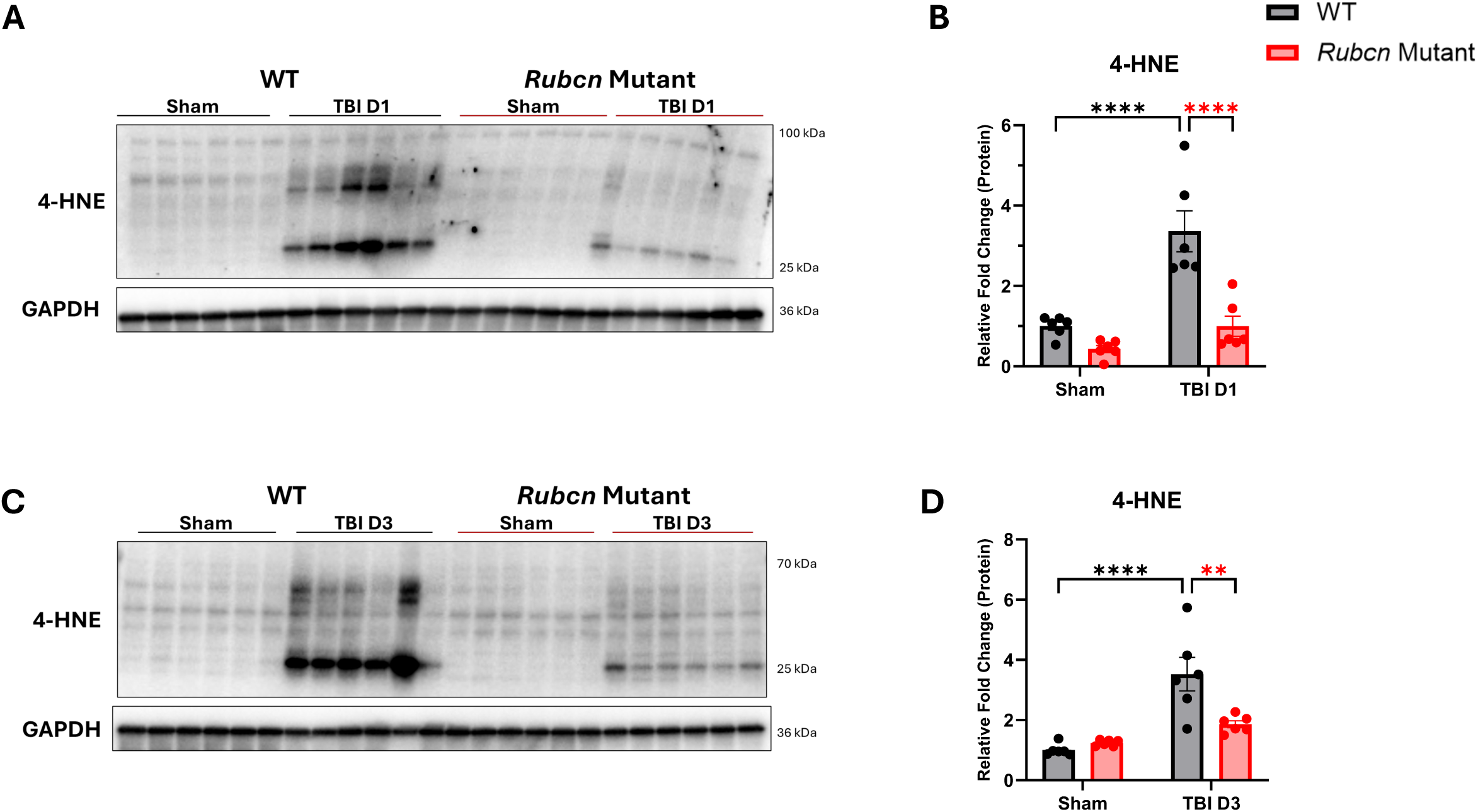
Rubicon mutation attenuates oxidative damage after TBI. **A** Immunoblot of ipsilateral cortical tissue lysates from wild-type and Rubcn-mutant mice comparing lipid peroxidation marker at 1 dpi. **B** Densitometric analysis of immunoblot in A. **C** Immunoblot of ipsilateral cortical tissue lysates from wild-type and Rubcn-mutant mice comparing lipid peroxidation marker at 3 dpi. **D** Densitometric analysis of immunoblot in C. Data represent mean ± s.e.m. Sample size (n) were as follows: WT Sham=6, WT TBI D1=6, WT TBI D3=6, Rubcn Mut. Sham=6, Rubcn Mut. D1=6, Rubcn Mut. D3=6.Statistical analyses: Two-way ANOVA with post hoc Tukey’s test and significance is represented by * P-value<0.05, ** P-value<0.005, *** P-value<0.0005, **** P-value<0.0001.

## Discussion

Rubicon has been implicated as an anti-inflammatory mediator through its role in CASM pathways including LAP and LANDO. In chronic neurodegenerative and some autoimmune conditions, Rubicon deficiency has been associated with exacerbated inflammation, impaired cargo clearance, and worsened pathology. (21, 23–25) However, Rubicon can also promote inflammation through negative regulation of autophagy. In ageing and metabolic diseases, Rubicon has been shown to suppress autophagy and mitophagy, promoting oxidative stress, proteotoxic burden, and inflammaging-like phenotypes. (16, 48–51) These dual roles indicate that the function and the mechanisms of Rubicon need to be individually evaluated in every experimental and disease context. Using a mouse model of TBI, we demonstrate that Rubicon mutation can attenuate acute inflammation and promote functional recovery after injury. Our findings indicate that in the context of brain injury Rubicon promotes inflammation by contributing to injury-induced autophagy inhibition.

We observed autophagy inhibition, elevated damage-associated markers indicative of cell damage, and upregulation of inflammatory responses in wild-type mice after injury. These changes were all attenuated in *Rubcn*-mutant mice. Impaired microglial and monocyte autophagy after brain injury has been shown by our lab to exacerbate inflammatory responses. (3, 4, 52) Consistent with this framework, reduced autophagy inhibition in Rubicon mutant mice correlated with diminished activation of inflammatory pathways. These data suggest that in acute pathological conditions, Rubicon-mediated suppression of canonical autophagy may dominate over its ability to promote CASM functions.

In addition to microglia and monocyte inflammatory polarization, hallmarks of TBI include neuronal cell death, which we also demonstrated to be affected by injury-induced inhibition of autophagy. (4) While we observed decrease in cell damage markers such as HMGB1 and α-fodrin in injured *Rubcn*-mutant mice, neither overall TBI-induced tissue loss nor levels of apoptosis (data not shown) were significantly altered between genotypes. Neuronal Rubicon has been reported to regulate mitophagy under mitochondrial stress conditions. (41) However, it is possible that under injury conditions, Rubicon can regulate microglial and monocyte but not neuronal autophagy. In that case, decrease in damage markers could be attributed to improved phagocytic removal of DAMPs or decrease in secondary inflammation-mediated damage. Alternatively, increased autophagy in *Rubcn*-mutant mice could attenuate injury in less damaged neurons but unable to protect severely damaged ones from death. Further experiments will be necessary to distinguish between these possibilities.

Our biochemical analyses reveal that the STOP codon present in the *Rubcn*-mutant mice does not result in a complete null but rather leads to production of a truncated RUBCN protein, lacking the N-terminal RUN domain while retaining downstream regions. Similar truncated mutant of Rubicon has been previously shown to enhance autophagy in B cells. (42) In our study, we saw baseline autophagy increase in Rubicon mutant macrophages and decreased inhibition of autophagy in mutant mice exposed to TBI. The truncated Rubicon protein in *Rubcn*-mutant mice is architecturally similar to Pacer, a positive regulator of autophagy, that shares PIKBD and RH domains. Pacer selectively recognizes phospho-Ser72 Rab7, activates VPS34, boosts PI3P production, recruits WIPI2 and ATG12/5/16L1 complex and promotes autophagy. (53) It is plausible that mutant Rubicon follows similar mechanisms, however, additional studies, including comparing full *Rubcn* knock-out to the N-terminally truncated mutant, will be necessary.

A mechanistic insight from our study is the identification of Rubicon as a regulator of injury-induced oxidative stress. Our data demonstrate that *Rubcn*-mutant mice exhibit significantly reduced lipid peroxidation following injury, indicating diminished ROS-mediated damage. Through unbiased proteomic analysis, we identify NRROS as a Rubicon interacting protein that preferentially associates with wild-type Rubicon. NRROS is known to limit NOX-2 activity by promoting degradation of the Gp91^phox^ sub-unit and suppressing phosphorylation of p47^phox^. (31–33) Since decreased interaction between RUBCN and NRROS in *Rubcn*-mutant mice were associated with decreased ROS-mediated damage, we expect that RUBCN inhibits NRROS function. One possible model could be that wild-type Rubicon sequesters NRROS following injury, limiting its ability to restrain NOX2 activity and thereby promoting ROS generation. In *Rubcn*-mutant mice, reduced interaction with RUBCN could increase NRROS function, resulting in decreased oxidative damage and attenuated inflammation. Decrease in ROS could also be an important factor contributing to decreased cell damage markers we observed in *Rubcn*-mutant mice after TBI.

Several limitations of this study warrant consideration. Our transcriptomic analyses were performed on bulk cortical tissue, which may mask cell-type specific effects of Rubicon mutation. Future studies employing cell-type specific genetic approaches, single cell transcriptomics and targeted manipulation of ROS signaling will be essential to fully delineate Rubicon’s multifaceted role in the injured brain. Additionally, while our interaction studies identify NRROS as Rubicon associating protein, the precise binding interface and regulatory dynamics of this interaction require further investigation. Finally, all our experiments were performed in male mice, and the applicability to females will need to be confirmed.

In summary, our findings reveal pro-inflammatory role for Rubicon in acute TBI, mediated through regulation of autophagy and oxidative stress. By linking Rubicon to NRROS-dependent control of ROS production, this study provides new mechanistic insight into how autophagy-associated proteins shape inflammatory outcomes after brain injury. These results underscore the importance of context-dependent assessment of Rubicon function and suggest that targeting Rubicon-mediated pathways may represent a novel therapeutic strategy to limit secondary injury and improve recovery after TBI.

## Methods

### Mouse studies

Rubicon mutant mice on C57BL/6 genetic background were obtained from the Jackson Laboratory (stock #032581). Transgenic GFP-LC3 mice on C57BL/6 genetic background were created by Dr. Noboru Mizushima and obtained from the Beth Levine group.(39) All mice were housed in cages with sterilized bedding at the University of Maryland, Baltimore (UMB) animal facility and maintained on a 12-hour light/dark cycle. GFP-LC3 (tg/+) Rubicon mutant mice were generated by crossing GFP-LC3 and Rubicon mutant mice at the UMB animal facility and were genotyped according to the Jackson Laboratory genotyping protocol. All *in vivo* experiments were performed on 12-16 weeks old male mice in accordance with the NIH guidelines for animal research and under an approved IACUC protocol at UMB. For tissue collection, mice were anesthetized and perfused transcardially with ice-cold saline. Ipsilateral and contralateral cortical and hippocampal tissues were harvested and flash-frozen until further processing for mRNA and protein analysis. *In vitro* experiments were performed using BMDMs derived from 12-16 weeks old male and female mice.

### Controlled Cortical Impact (CCI)

Mice were subjected to moderate TBI using a CCI device customized with a microprocessor-regulated pneumatic impactor equipped with a 2.3-mm diameter tip as described previously. (3, 4, 52) Briefly, mice were anesthetized with 3% isoflurane and underwent a 4-mm craniotomy over the left parietal bone following a 10-mm midline scalp incision. Moderate CCI was induced on the exposed portion of the brain using the pneumatic impactor at an impact velocity of 4 m/s, a deformation depth of 1.5 mm, and a dwell time of 100 ms. The incision sites were then closed with sterile sutures. Sham mice were subjected to the same procedures, excluding the craniotomy and impact.

### Neurobehavioral assessment

**Beam Walk**: To evaluate fine motor coordination, beam walk test was performed at several time points after CCI induction. (3, 54) Mice were trained to walk on a wooden beam (approximately 5 mm and 120 cm long) for 3 consecutive days prior to injury. Testing was conducted on 1,3, 7, 14, 21, and 28 dpi. The number of foot faults for the right hind-limb was recorded out of 50 total steps taken on the beam.

**CatWalk:** To evaluate motor function and gait, analysis was performed at multiple time points using the CatWalk XT (Noldus) automated system as described previously. (55) Mice traversed the CatWalk apparatus which records paw print position to analyze gait posture and weight distribution. Each mouse completed a minimum of three valid runs, defined as continuous crossings without turning or side-wall scaling. Non-compliant runs were omitted from the analysis. One-two foot-support was calculated by summing the percentages of Single-, Diagonal-, Girdle-, and Lateral-Support. Three-Four foot-support was calculated by summing the percentages of Three– and Four-Support.

### RNA-seq and analyses

Total RNA was extracted from the frozen ipsilateral cortical tissues using QIAzol Lysis reagent (Qiagen; 79306) and purified with the RNeasy mini kit (Qiagen; 74104) according to the manufacturer’s protocol. Bulk RNA sequencing was performed at the University of Maryland Institute for Genome Sciences.

**Galaxy and Metabolanalyst:** Differentially expressed genes (DEGs) were generated using the DESeq2 tool in the Galaxy web platform (usegalaxy.org, version 2.11.40.8_galaxy0). (56, 57) Variant sparse partial least squares discriminant analysis (sPLS-DA) plot and heat maps were created in Metaboanalyst 6.0 using data from the DESeq2 normalized counts. (58) Volcano plots were generated in Galaxy with the following parameters: *P*-adj<0.05; log_2_(fold-change)>1 for TBI D3 over Sham samples; *P*-adj<0.05; log_2_(fold-change)>0.25 for *Rubcn* mutant over wild-type samples.

**DAVID Analysis:** The *Rubcn*-mutant vs. wild-type TBI D3 DEGs generated from Galaxy were filtered for significance (*P*-value<0.01) and sorted as upregulated or downregulated based on their log_2_(fold-change) values. The resulting upregulated and downregulated DEGs were uploaded separately into the DAVID functional annotation tool.(59) Bar graphs were created using Gene Ontology Biological Process Direct (GO:BP) terms sorted by FDR (<0.05).

**Ingenuity Pathway Analysis (IPA):** Differential expression results comparing the effect of injury in *Rubcn*-mutant mice (TBI D3 by Sham; *P*-value <0.002) to the effect of injury in wild-type mice (TBI D3 by Sham; *P*-value <0.002) were analyzed via IPA (QIAGEN) using canonical pathway enrichment and upstream regulator prediction modules.

### Quantitative real-timePCR (qRT-PCR)

Total RNA was extracted from the frozen cortices as described above. Two μg of RNA were reverse-transcribed using the High-Capacity cDNA Reverse Transcription Kit (Applied Biosystems; 4374967). Real-time quantitative PCR was performed using the TaqMan™ Universal Master Mix II (Applied Biosystems; 4440040) and mouse TaqMan Gene Expression Assays on a QuantStudio^TM^ System (ThermoFisher Scientific). The following Assay IDs were used for mRNA quantification: Mm00434228_m1(*Il1b*), Mm00524817_m1 (*Nrros*), Mm99999915_g1 (*Gapdh*), Mm01307193_g1 (*Apoe*), Mm00442646_m1 (*Abca1*), Mm00443451_m1 (*Nr1h3*), Mm00437390_m1 (*Abcg1*), Mm00840904_m1 (*Nlrp3*), Mm00438023_m1 (*Casp1*), Mm01158117_m1 (*Tmem173*), Mm01147496_m1(*Mb21d1*), Mm01184322_m1 (*Pparg*), Mm00476361_m1 (*Nfkb1*), Mm00448091_m1 (*Sqstm1*), Mm00782868_sH (*Map1lc3b*), Mm01324470_m1 (*lcn2*), Mm01178820_m1 (*Tgfb1*). *Gapdh* was used as a normalization control.

### Immunoblotting

Ipsilateral cortical and hippocampal tissues were homogenized in cold radioimmunoprecipitation assay (RIPA) buffer (Teknova; R3792) containing protease inhibitor (Roche; 11836170001) and phosphatase inhibitors (SigmaAldrich; P5726; P0044) and centrifuged as described previously. (3, 4) The resulting supernatants were used to measure protein concentrations using a bicinchoninic acid (BCA) assay (ThermoFischer Scientific; 23225). Samples were combined with 2x Laemmli buffer, separated by 4-20% sodium dodecyl sulfate-polyacrylamide gel (Bio-Rad; 5671095) electrophoresis and transferred onto polyvinylidene difluoride membranes (Bio-Rad; 12023927) using a semi-dry transfer system. Membranes were blocked with 5% non-fat milk in Tris-buffered saline with 0.1% Tween 20 (TBS-t) for 1 h at room temperature (RT). Membranes were then incubated with primary antibodies in 5% BSA in TBS-t overnight at 4 °C, followed by incubation with HRP-conjugated secondary antibodies in 5% non-fat milk in TBS-t for 1 h at RT. Membrane blots were developed using SuperSignal West Pico (ThermoFischer Scientific; 34577), SuperSignal West Femto (ThermoFischer Scientific; 34095), or SuperSignal West Atto (A38554), and chemiluminescent signals were detected using a Universal Hood II Gel Doc system (Bio-Rad). Protein signals were quantified using Image Lab software (Bio-Rad Laboratories, Inc., Hercules, CA; version 6.1.0 build 7) and normalized to loading controls. The following antibodies were used for immunobloting: RUBCN (CST-8465S; 68261S), NRROS (CST; 34388S), NLRP3 (CST; 15101S), cGAS (CST; 15102S), STING (CST; 13647S), α-fodrin (BML-FG6090-0100), P62 (BD Biosciences; 610832), LC3(CST; 2775S; NB100-2220), GAPDH (CST; 2118S), HMGB1(CST; 3935S), FLAG (CST; 33542S), MYC (CST; 2276S), HIS (CST; 2365S), GFP (CST; 2955S), 4-HNE (R&D Systems; MAB3249), mouse IgG1(CST; 5415S), mouse IgG2a (CST; 61656), anti-rabbit IgG-HRP (CST; 7074S), and anti-mouse IgG-HRP (CST; 7076S) Uncropped immunoblots are presented if **Fig. S5**.

### Immunohistochemistry and image acquisition/analyses

Mice were anesthetized and perfused transcardially with ice-cold saline and then with ice-cold 4% paraformaldehyde. Whole brains were collected and post-fixed in 4% paraformaldehyde overnight at 4 °C. The samples were cryoprotected in 20% and 30% sucrose in PBS subsequently at 4 °C overnight and sectioned into 30 μm or 60 μm thick frozen coronal slices. The sections were stored in section storage solution (FD NeuroTechnologies; PC101) for free-floating immunofluorescence (IF) staining or mounted directly onto glass slides for histology.

For IF staining, frozen sections were allowed to thaw to RT and rinsed with PBS. The sections were incubated in 100 mM Glycine (in PBS) and permeabilization buffer (0.5% Triton X-100 in PBS) for 30 min each with orbital rotation. They were then blocked in blocking buffer (5% normal goat serum, 1% BSA, 0.1% Triton X-100 in PBS) for 1 h at RT and incubated with primary antibodies in blocking buffer overnight at 4 °C. Sections were incubated with secondary antibodies in blocking buffer for 2 h at RT and counterstained with Hoechst (Invitrogen; H1399). The sections were mounted on glass slides and covered with a thin layer of Hydromount solution (National Diagnostics; 5089990144) and a glass cover slip. The following antibodies were used: IBA1 (FujiFilm Wako; 019-19741), SQSTM1/P62 (Progen; GP62-C), GFAP (Dako; Z0334), goat α-guinea pig Alexa Fluor 568 (Invitrogen; A-11075), and goat α-rabbit Alexa Fluor 633 (Invitrogen; A-21070).

Immunofluorescent images of the peri-lesion area were acquired on a Nikon Eclipse Ni-E microscope, with emission wavelengths including DAPI (420-460 nm), FITC (515-555 nm), Cy3 (577-630 nm) and Cy5 (677-711 nm), using 10x and 20x objective lenses. Constant exposure times were maintained for all sections within the experiment. Analysis of cortical volume loss following CCI was performed using serial coronal sections (60 μm thick) collected at 180 μm intervals as described before.(60) Cortical area was quantified in NeuN stained sections by tracing cortical boundaries in each hemisphere using Fiji (ImageJ 1.54f). (61) The cortical region was defined as the area between the dorsal aspect of the corpus callosum and the pial surface. The percent cortex remaining was calculated as the ratio of the summed ipsilateral cortical area to the summed contralateral cortical area across sections. Representative images for GFP-LC3 coronal sections stained with P62 and IBA were deconvolved using TRUESHARP online deconvolution (Version 1.0, Abberior Instruments GmbH, Gӧttingen, Germany; https://app.truesharp.rocks/). (62) All analyses were performed on raw images. P62 positive cells within IBA/LC3 positive cells were analyzed using NIS-Elements software (Nikon; V5.30.06) as follows: nuclei were identified using Bright Spot Detection parameter, while IF positive cells were detected using Detect Regional Maxima and global Threshold parameters. Data were normalized to the total number of nuclei per image.

### Primary macrophage culture and autophagic activity assay

Bone marrow derived macrophages (BMDMs) were differentiated from primary cells using Roswell Park Memorial Institute (RPMI) 1640 Medium (Gibco; 22400105), supplemented with 10% fetal bovine serum (FBS) (GeminiBio; 900-108H-500), 10% L929-conditioned medium, and 1% antibiotic-antimycotic (Gibco; 15240062), as described before. (52) Briefly, bone marrow was flushed from the femora and tibiae of mice using ice-cold PBS. The marrow was centrifuged into form a pellet and suspended in ice-cold RBC lysis buffer for 5-10 minutes. The cell suspension was neutralized with warm BMDM medium and centrifuged again to pellet the cells. The cell pellet was resuspended in BMDM medium, filtered through 70 μm cell strainers, and cultured in T75 flasks. Primary cells were allowed to differentiate into BMDMs for 7 days, with medium changes every 2-3 days. For the autophagic activity assay, BMDMs were harvested on day 7 and re-seeded into 12 well plates. Cells were pre-treated with 50 nM bafilomycin A1 (CST; 54645S) for 1 h and serum-starved for 4 h in the presence of bafilomycin A1 treatment. Cells were washed with ice-cold PBS before being lysed with RIPA buffer for immunoblotting. Immunoblot analysis was performed as described above.

### Immunoprecipitation of endogenous protein

Following saline perfusion, whole brain tissues were removed and homogenized in cold radioimmunoprecipitation assay (RIPA) buffer (Teknova; R3792) containing protease inhibitor (Roche; 11836170001) and phosphatase inhibitors (SigmaAldrich; P5726; P0044) using a glass homogenizer. Samples were allowed to lyse further for 30 min at 4 °C with rotation and then centrifuged. Protein concentrations of the resulting supernatants were measured using BCA assay, and equal amounts of protein were pre-cleared with Pierce^TM^ Protein A/G Agarose beads (ThermoFisher Scientific; 20421) for 2 h at 4°C with rotation. Pre-cleared samples were incubated with the indicated primary antibodies overnight at 4°C and then with agarose beads for 2 h the following day at 4°C with rotation. The protein-agarose bead complexes were washed with RIPA buffer four times and stored at –80 °C until processed for mass spectrometry analysis. For immunoblot analysis, proteins were eluted from the beads in 2x Laemmli buffer by boiling for 5 min at 95 °C.

### Mass spectrometry-based proteomics

Mouse brain lysates were immunoprecipitated using Rubicon antibodies (Rb-A and Rb-B) and stored at −80 °C until further analysis. Cell lysis and protein digestion were performed as previously described. (63–65) Briefly, samples were lysed in a lysis buffer containing 5% sodium dodecyl sulfate (Sigma-Aldrich; L4509) and 50 mM triethylammonium bicarbonate (1 M, pH 8.0) (Sigma-Aldrich; 7408). Proteins were extracted and digested using S-trap micro columns (ProtiFi, NY; C002-MICRO). The eluted peptides from the S-trap column were dried, and peptide concentration was determined using BCA assay (ThermoFischer Scientific; 23225) after reconstitution in 0.1% formic acid. All tryptic peptides were separated on a nanoACQUITY Ultra-Performance Liquid Chromatography analytical column (BEH130 C18, 1.7 µm, 75 µm x 200 mm; Waters Corporation, Milford, MA, USA) over a 185 min linear acetonitrile gradient (3–40%) with 0.1% formic acid on a nanoACQUITY Ultra-Performance Liquid Chromatography system (Waters Corporation, Milford, MA USA) and analyzed on a coupled Orbitrap Fusion Lumos Tribrid mass spectrometer (Thermo Scientific, San Jose, CA USA). Full scans were acquired at a resolution of 240,000, and precursors were selected for fragmentation by high-energy collisional dissociation of 35% for a maximum 3 s cycle. The MS/MS raw files were processed using Proteome Discoverer (PD, version 2.5.0.400; Thermo Fisher Scientific) with the Sequest HT search engine against the UniProt mouse reference proteome (release 2022.06, 17,096 entries). Searches were configured with static modifications for carbamidomethyl on cysteines (+57.021 Da), dynamic modifications for the oxidation of methionine residues (+15.995 Da), precursor mass tolerance of 20 ppm, fragment mass tolerance of 0.5 Da. Trypsin was used as a digestion enzyme with a maximum of two missed cleavages. The minimum and maximum peptide lengths were set to 6 and 144, respectively. Label-free quantification was performed using the Minora feature detector, a tool embedded in the PD bioinformatics platform.(66) For high-confidence results, protein identification was filtered to a 1% false discovery rate (FDR) in peptide spectra match (PSM), peptide, and protein levels. The FDR was calculated using the Percolator algorithm embedded in PD. Next, the exported protein abundance values were analyzed and visualized using Perseus software (version 1.6.14.0). To ensure high confidence in statistical analysis, data were further filtered to include only proteins identified without missing values across biological samples. The quantitative protein data were log_2_ transformed and normalized using median centering. Comparative enrichment analysis was performed by normalizing the abundance of each group immunoprecipitated with each RUBCN antibody to the average abundance in the IgG immunoprecipitates. The fold change for enrichment in wild-type samples was calculated for each antibody following normalization.

### Immunoprecipitation of exogenous protein

HEK293T cells were maintained in Dulbecco’s Modified Eagle Medium (DMEM) (Gibco; 11995073) supplemented with 10% fetal bovine serum (FBS) (GeminiBio; 900-108H-500) and 1% antibiotic-antimycotic (Gibco; 15240062). HEK293T cells were transiently transfected with EGFP-Rubicon (Addgene; 28022), EGFP-Rubicon Δ182 (Addgene; 28041), EGFP-Rubicon ΔCT (Addgene; 28040), pcDNA3.1(+)-human LRRC33 (Addgene; 111602), and/or pEGFP-C1 vectors using Lipofectamine^TM^ 3000 Transfection Reagent (ThermoFisher Scientific; L3000015) for 24 h.(12, 44, 46) The cell monolayers were washed with ice-cold PBS and lysed in cell lysis buffer (50 mM Tris-HCl, 150 mM NaCl, 1 mM EDTA, and 1% Triton X-100 or 0.5% Igepal CA-630 (SigmaAldrich; I8896) containing protease and phosphatase inhibitors. The cell lysates were centrifuged at 18,000 xg for 30 min, and the resulting supernatants were pre-cleared with Pierce^TM^ Protein A/G Agarose beads (ThermoFisher Scientific; 20421) and mouse mAb IgG1 (CST; 5415) or mouse IgG2a (CST; 61656) for 2 h at 4 °C with rotation. For FLAG pull-down, pre-cleared supernatants were then incubated with Anti-FLAG M2 Affinity Gel (SigmaAldrich; A2220) or Anti-HA Agarose clone HA-7(SigmaAldrich; A2095) for another 2 h at 4°C with rotation. For MYC pull-down, pre-cleared supernatants were incubated with MYC (CST; 2276S) or mouse IgG2a overnight, followed by incubation with agarose beads for 2 h at 4 °C with rotation. Following incubation, the beads were washed three times with ice-cold wash buffer (50 mM Tris-HCl, 150 mM NaCl). Proteins were eluted from the beads with 2x Laemmli buffer, boiled at 95 °C for 3 min, and analyzed by immunoblotting.

### Statistical analyses

Sample sizes for animal experiments are consistent with power analyses (power 80-90%, alpha 0.05) based on experience with the CCI mouse model and expected effect sizes within the TBI field. Animals were block randomized into injury vs sham groups. Data are presented as mean ± s.e.m. Statistical analyses were performed using GraphPad Prism (v 10.1.0 (316); GraphPad Software, Boston, MA). Unless otherwise indicated, grouped datasets were analyzed by two-way ANOVA followed by Tukey’s multiple comparisons test. Statistical significance was defined as *P*-value<0.05. Sample sizes and specific analyses for each experiment are detailed in the figure legends. Behavior data were analyzed using mixed-effects analysis with a post hoc Tukey’s test. An unpaired Student’s t-test was used to assess the significance of the percentage of cortical damage. Protein signal from immunoprecipitation experiment was analyzed using one-way ANOVA followed by Tukey’s post hoc test.

## Acknowledgements

We thank Dr. Bradley Heckmann (University of South Florida) for advice on the study of Rubicon function.

## Author Contributions

The project was conceived by MML; experiments were designed by ST, CS, MAK and MML; experiments were performed by ST, AM, OP, DPHN, MMW, CS and MK; data were analyzed by ST, MMW, MAK and MML; the manuscript was written by ST and MML with comments from all authors.

## Data Availability Statement

All processed data is included with the manuscript files; raw data will be available upon publication.

